# Non-physiologic interaction of MG53 with insulin receptor and lack of evidence for MG53’s role in controlling insulin-stimulated Akt phosphorylation in muscle, heart and liver tissues

**DOI:** 10.1101/2024.01.26.576909

**Authors:** Kyung Eun Lee, Miyuki Nishi, Jongsoo Kim, Takashi Murayama, Zachary Dawson, Xiaoliang Wang, Xinyu Zhou, Tao Tan, Chuanxi Cai, Hiroshi Takeshima, Ki Ho Park

## Abstract

**Rationale:** MG53’s known function in facilitating tissue repair and anti-inflammation has broad applications to regenerative medicine. There is controversy regarding MG53’s role in the development of type 2 diabetes mellitus (T2DM).

**Objective:** This study aims to address this controversy – whether MG53’s myokine function contributes to inhibition of insulin signaling in muscle, heart, and liver tissues.

**Study Design:** We determined the binding affinity of the recombinant human MG53 (rhMG53) to the insulin receptor extracellular domain (IR-ECD) and found low affinity of interaction with K_d_ (>480 nM). Using cultured C2C12 myotubes and HepG2 cells, we found no effect of rhMG53 on insulin-stimulated Akt phosphorylation (p-Akt). We performed in vivo assay with C57BL/6J mice subjected to insulin stimulation (1 U/kg, intraperitoneal injection) and observed no effect of rhMG53 on insulin-stimulated p-Akt in muscle, heart and liver tissues.

**Conclusion:** Overall, our data suggest that rhMG53 can bind to the IR-ECD, however has a low likelihood of a physiologic role, as the K_d_ for binding is ∼10,000 higher than the physiologic level of MG53 present in the serum of rodents and humans (∼10 pM). Our findings question the notion proposed by Xiao and colleagues – whether targeting circulating MG53 opens a new therapeutic avenue for T2DM and its complications.

## RESULTS AND DISCUSSION

Since the discovery of MG53 (TRIM72) as a cell membrane repair protein in 2009^1^, notable progress has been made in understanding its mechanistic action in regenerative medicine and in regulation of metabolic syndromes. In addition to facilitating tissue repair, MG53 also has anti-inflammation function with broad applications to aging biology, organ failure, viral infection and wound healing^2^. However, MG53’s role in the development of type 2 diabetes mellitus (T2DM) continues to be debated.

Song et al^3^ postulate that MG53-mediated downregulation of IRS-1 serves as a causative factor for T2DM, while other experts in the field challenge their conclusion^4^. Wu et al^5^ reported elevated serum levels of MG53 (∼2 fold) in diabetic mouse models and human diabetic patients, concluding that MG53 has dual actions as a myokine and an E3 ligase to inhibit insulin signaling in muscle, heart and liver tissues. Using BiACore assay, they showed recombinant human MG53 (rhMG53) protein binds to the extracellular domain of insulin receptor (IR-ECD) to block insulin signaling, with an estimated K_d_ of 8 nM (Figure 6C Wu et al^5^). Compared to the rapid k_off_/k_on_ of insulin binding, the kinetics for rhMG53 binding to IR-ECD is atypical; rhMG53 has slow reaction and fails to reach equilibration or saturation (Figure 6B Wu et al^5^). We have followed the exact protocol as described and found that rhMG53 can interact with IR-ECD but at a much higher K_d_ (>480 nM) (**Figure A** and **B**). Moreover, we confirmed pre-loading of the Octet sensorgrams with rhMG53 does not impact insulin’s affinity with ECD (data not shown). Our results indicate that rhMG53 does not directly interfere with insulin binding to IR-ECD, and rhMG53 has a much lower competitive/allosteric threshold than previously described.

Wu et al^5^ further suggested that “… extracellular MG53 acts as an allosteric, rather than a competitive, blocker of the IR”, based on their findings in liver cell culture where incubation with rhMG53 (not BSA) inhibited insulin-stimulated phosphorylation of Akt (p-Akt) (Figure 7A Wu et al^5^); and intravenous administration of rhMG53 blocked insulin-stimulated p-Akt in mouse tissues (Figure 5A Wu et al^5^). We were puzzled by these results, as a recent report by Philouze et al^4^ showed negative findings with rhMG53 on insulin-stimulated p-Akt (Figure 3 and 4 Philouze et al^4^). Thus, we conducted a study to determine whether rhMG53 alters insulin-stimulated p-Akt in a dose-dependent manner. We treated C2C12 myotubes and HepG2 cells with 10 nM of insulin, then quantified the impact of rhMG53 in a wide range of concentrations (10, 100 and 500 nM). As shown in **Figure C** and **D**, we saw no effect of rhMG53 on insulin-stimulated p-Akt in all concentrations tested.

These findings prompted us to conduct *in vivo* assays to determine whether intravenous administration of rhMG53 has any impact on insulin-stimulated p-Akt in muscle, heart and liver tissues. Following the protocol of Wu et al^5^, C57BL6J mice (male, 8-10 weeks old, The Jackson Laboratory) were treated with 1 U/kg of insulin (intraperitoneal injection); 10 min prior to insulin injection, 1 mg/kg or 6 mg/kg of rhMG53 was administered into the tail vein of the mice. Administration of 1 mg/kg rhMG53 would result in a peak level of MG53 in the blood (∼10 µg/ml) which should be 1000-fold and 500-fold higher than the resting level of MG53 observed in healthy and diabetic animals. We would also expect this to further saturate the IR-ECD if any allosteric interaction was occurring. However, we were surprised to find that rhMG53 has no impact on p-Akt level in skeletal muscle, heart or liver tissues (**Figure E**), which contrasts the data presented by Wu et al, who notably used 6 mg/kg of rhMG53 in their study.

Overall, our data suggest that rhMG53 can bind to IR-ECD, however has low likelihood of a physiologic role, as the K_d_ for binding is ∼10,000 higher than the physiologic level of MG53 present in the serum of rodents and humans (∼10 pM). Consistent with the data shown by Wu et al^5^, rhMG53 clearly does not interfere with insulin binding at the IR. Based on our current findings and those of Philouze et al^4^, it is unlikely that MG53 has any allosteric impact on insulin-signaling in muscle, heart and liver tissues. All together, these findings question the notion proposed by Wu et al^5^ – whether targeting circulating MG53 opens a new therapeutic avenue for T2DM and its complications.

## MATERIAL AND METHODS

### Cell culture

Cell lines were maintained at 37 °C, 95 % air, and 5 % CO_2_ in a humidified incubator. C2C12 cells (ATCC, CRL-1772) were cultured in Dulbecco’s Modified Eagle’s Medium with 10 % fetal bovine serum (FBS, Gibco^TM^, 16140071) and 1 % penicillin-streptomycin (Gibco^TM^, 15140122). For differentiation, C2C12 cells were cultured in the DMEM with 2.5% horse serum (Gibco^TM^, 26050088) for 5 days. HepG2 cells (ATCC, HB-8065) were cultured in RPMI1640 medium with 10 % FBS and 1 % penicillin-streptomycin.

### Animal Care and Experimental Mice

Animal handling and experimental procedures were performed according to protocols approved by the Institutional Animal Care and Use Committee (IACUC) of University of Virginia and were compliant with guidelines of the American Association for the Accreditation of Laboratory Animal Care. All experimental mice purchased from The Jackson Laboratory were 8-10 weeks age (C57BL/6J, The Jackson Lab stock No: 000664).

### Treatment of rhMG53 and insulin

HepG2 and differentiated C2C12 cells were pre-treated with 10, 100, or 500 nM of rhMG53 (or 500 nM of BSA as control) for 60 minutes, and then stimulated with 10 nM of insulin (Gibco^TM^, 12585014) for 10 minutes. Cells were washed using ice-cold PBS and detached mechanically using a scraper, and then collected.

For *in vivo* treatment, after fasting overnight (13-15 hours), mice were administered rhMG53 and BSA intravenously at doses of 1 mg/kg and 6 mg/kg, respectively. 10 minutes later, they were subjected to stimulation with 1 U/kg of insulin via intraperitoneal injection. The heart, liver, and skeletal muscles were collected 10 minutes after insulin stimulation, and snap-frozen in liquid nitrogen, and stored in -80 °C for later processing and analysis.

### Immunoblotting

Whole lysates were isolated from cells (C2C12 and HepG2) and mouse tissues (heart, liver, and skeletal muscles) with RIPA lysis buffer containing protease inhibitor (Sigma, P8340) and phosphatase inhibitors (Sigma, P0044). The denatured proteins (20 µg/well) were separated by 4-12 % protein gel (Thermo Scientific) and transferred onto PVDF membranes (MiliporeSigma, IPVH00010). Membranes were incubated in 5% (w/v) nonfat dry milk for 1 hour at room temperature, further probed with primary antibody and incubate at 4 °C with gentle shaking overnight. Then, they were washed with Tris-buffered saline with 0.1 % Tween 20 detergent (TBST) and probed with a HRP (horseradish peroxidase)-conjugated secondary antibody. Immunodetection was performed by iBright^TM^ CL 1500 Imaging System (Invitrogen) using SuperSignal West Pico PLUS Chemiluminescent Substrate (Thermo Scientific, 34580). Primary antibodies used were as follows: Akt (Cell Signaling Technology, 9272S, 1:1000 dilute), phospho-Akt (Ser 473) (Cell Signaling Technology, 4060S, 1:2000 dilute), GAPDH (Cell Signaling Technology, 2118S, 1:5000 dilute) was diluted in 2.5 % (w/v) nonfat dry milk.

### Biolayer Interferometry (BLI) Assay

The binding affinity of rhMG53 with human insulin receptor extracellular domain (IR ECD)-His (INR-H52Ha, Acro Biosystems) was determined using Anti-Penta-His (HIS1K) biosensors in an Octet Red 96 instrument (ForteBio Inc., Menlo Park, CA). IR ECD-His was immobilized onto the surface of HIS1K biosensors. Increasing concentrations of rhMG53 and CNP were allowed to interact with the immobilized IR ECD at room temperature in running buffer (10 mM Hepes, 150 mM NaCl, 0.005% Tween-20 pH 7.4). The final volume of all solutions was 200 µL. Assays were performed in black solid 96-well flat bottom plates on a shaker set at 1,000 rpm/min. The association and dissociation of IR ECD was measured for 250 sec interval. All data were analyzed using the Octet Data Analysis 9.0 software (ForteBio).

### Statistical analysis

ANOVA test (single factor) was used to determine statistical difference between groups. All data were analyzed using Excel and GraphPad Prism 10 software.

---

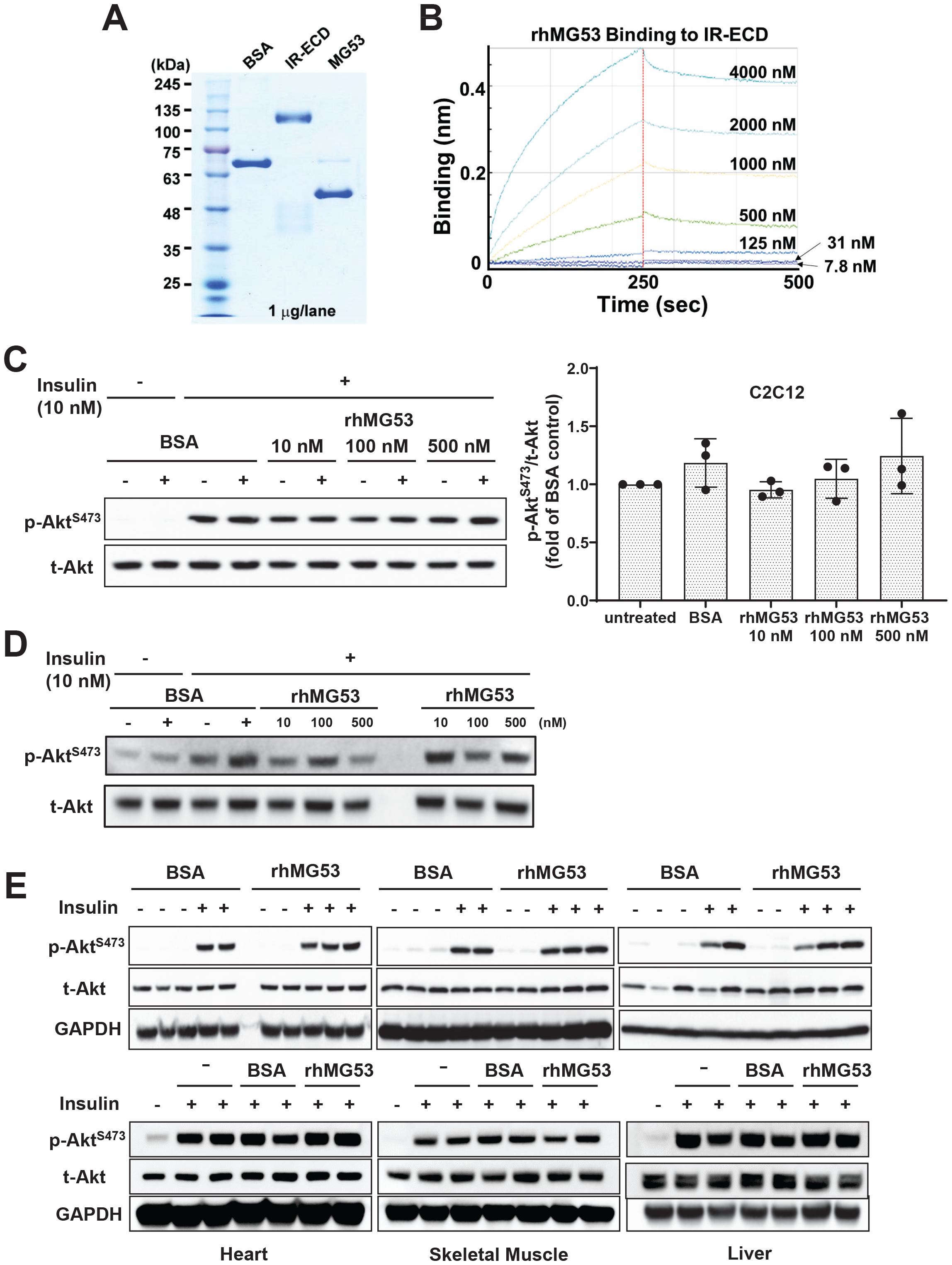
**(A)** SDS-polyacrylamide gel electrophoresis (SDS-PAGE) of bovine serum albumin (BSA, Sigma A1470), IR-ECD (Acro Biosystem), and rhMG53 (see ref 2 for preparation). **(B)** In the protein-protein interaction assay using Octet RED96 instrument (ForteBio Inc., Menlo Park, CA), Octet sensorgrams of rhMG53 binding to IR-ECD revealed a K_d_ > 480 nM (n=3). **(C)** C2C12 myotubes (5 days post differentiation) and **(D)** HepG2 cells were treated with varying concentrations of rhMG53 for 1 hour (or BSA as control), followed by 10 nM insulin for 10 min. Western blot were conducted with antibody against p-Akt (Cat #4060, Cell Signaling Technology (CST)) and total Akt (t-Akt) (Cat #9272, CST). **(E)** 1 mg/kg rhMG53 (upper panel) and 6 mg/kg rhMG53 (lower panel) treatment did not alter the level of p-Akt in skeletal muscle, heart and liver derived from mice subjected to 1 U/kg insulin treatment.

## References

1. Cai C, Masumiya H, Weisleder N, Matsuda N, Nishi M, Hwang M, Ko JK, Lin P, Thornton A, Zhao X, Pan Z, Komazaki S, Brotto M, Takeshima H, Ma J. MG53 nucleates assembly of cell membrane repair machinery. Nature cell biology. 2009;11:56–64.

2. Whitson BA, Tan T, Gong N, Zhu H, Ma J. Muscle multiorgan crosstalk with MG53 as a myokine for tissue repair and regeneration. Curr Opin Pharmacol. 2021;59:26–32.

3. Song R, Peng W, Zhang Y, Lv F, Wu HK, Guo J, Cao Y, Pi Y, Zhang X, Jin L, Zhang M, Jiang P, Liu F, Meng S, Zhang X, Jiang P, Cao CM, Xiao RP. Central role of E3 ubiquitin ligase MG53 in insulin resistance and metabolic disorders. Nature. 2013;494:375–379.

4. Philouze C, Turban S, Cremers B, Caliez A, Lamarche G, Bernard C, Provost N, Delerive P. MG53 is not a critical regulator of insulin signaling pathway in skeletal muscle. PLoS One. 2021;16:e0245179.

5. Wu HK, Zhang Y, Cao CM, Hu X, Fang M, Yao Y, Jin L, Chen G, Jiang P, Zhang S, Song R, Peng W, Liu F, Guo J, Tang L, He Y, Shan D, Huang J, Zhou Z, Wang D, Lv F, Xiao RP. Glucose-sensitive Myokine/Cardiokine MG53 Regulates Systemic Insulin Response and Metabolic Homeostasis. Circulation. 2018.

